# Altered Sensorimotor-to-Transmodal Hierarchical Organization in Schizophrenia

**DOI:** 10.1101/2020.03.06.980607

**Authors:** Debo Dong, Dezhong Yao, Yulin Wang, Seok-Jun Hong, Sarah Genon, Fei Xin, Kyesam Jung, Hui He, Xuebin Chang, Mingjun Duan, Boris Bernhardt, Daniel S. Margulies, Jorge Sepulcre, Simon B. Eickhoff, Cheng Luo

## Abstract

For decades, schizophrenia has been primarily conceptualized as a disorder of high-order cognitive functions with deficits in executive brain regions. Yet due to the increasing reports of early sensory processing deficit, recent models focus more on the developmental effects of impaired sensory process on high-order functions. The present study examined whether this pathological interaction relates to an overarching system-level imbalance, specifically a disruption in macroscale hierarchy affecting integration and segregation of unimodal and transmodal networks. We applied a novel combination of connectome gradient and stepwise connectivity analysis to resting-state functional magnetic resonance imaging (rsfMRI) to characterize the sensorimotor-to-transmodal cortical hierarchy organization (96 patients vs. 122 controls). Using these techniques, we demonstrated compression of the cortical hierarchy organization in schizophrenia, with a prominent compression from the sensorimotor region and a less prominent compression from the frontal-parietal region, resulting in a diminished separation between sensory and fronto-parietal cognitive systems. Further analyses suggested reduced differentiation related to atypical functional connectome transition from unimodal to transmodal brain areas. Specifically, we found hypo-connectivity within unimodal regions and hyper-connectivity between unimodal regions and frontoparietal and ventral attention regions along the classical sensation-to-cognition continuum established in prior neuroanatomical work. The compression of cortical hierarchy organization represents a novel and integrative system-level substrate underlying the pathological interaction of early sensory and cognitive function in schizophrenia. This abnormal cortical hierarchy organization suggests cascaded impairments stemming from the disrupted somatosensory-motor system and inefficient integration of bottom-up sensory information with attentional demands and executive control processes partially account for high-level cognitive deficits characteristic of schizophrenia.

## 1. Introduction

When “dementia praecox” was first proposed to describe schizophrenia by Kraeplin in the late 19^th^ century, cognitive deficit was regarded as the core component of the disorder (Dondé et al., 2019a), as evidenced by impaired function of higher-order brain regions, such as the prefrontal cortex (Minzenberg et al., 2009). Since then, perceptual deficits in the development of schizophrenia have been relatively ignored in the research field compared to the focus on the cognitive deficits. Although early sensory deficits have been well described in the schizophrenia literature, they have often been attributed to failures of attention and other top-down mechanisms and not been emphasized within prevailing psychiatric models (Rassovsky et al., 2005; Van Der Stelt et al., 2004). However, increasing empirical evidence has begun to highlight the importance of sensory processing dysfunction by ‘bottom-up’ dysregulation towards higher cognitive function in schizophrenia (Butler et al., 2007; Calderone et al., 2013; Dias et al., 2011; Dondé et al., 2019b; Hoptman et al., 2018; Leitman et al., 2010). Accordingly, recent models are gradually shifting the research focus more to the impaired developmental interactions between early sensory and high-order processes in schizophrenia (Javitt, 2009a; Javitt and Freedman, 2015; Javitt and Sweet, 2015). As such, the identification of neural mechanisms underlying the impaired functional integration between and within early sensory and cognitive brain systems is crucial to understand the pathophysiology mechanisms of schizophrenia, which ultimately helps guide future interventional approaches.

The aforementioned functional interaction can be explored by characterizing functional connectivity within and between different brain systems (van den Heuvel and Pol, 2010). Using resting-state functional connectivity (rsFC), researchers are able to observe abnormal FC within the high-order default, frontoparietal network as well as ventral attention network in schizophrenia (Dong et al., 2018b; Jiang et al., 2019; Liao et al., 2019; Pettersson-Yeo et al., 2011; Pläschke et al., 2017). Aside from these abnormalities, an increasing number of rsFC studies suggest dysfunctional intrinsic connectivity within visual and somatosensory systems in this condition (Bordier et al., 2018; Chen et al., 2016, 2014; Dong et al., 2018a; Guo et al., 2014; Jiang et al., 2015; Liu et al., 2018; Yiwen Zhang et al., 2019). Up to this point, only a few studies have looked at how sensory networks pathologically interact with higher-order association systems in schizophrenia (Berman et al., 2017; Hoptman et al., 2018; Kaufmann et al., 2015). In this more holistic view, brain dysfunction in schizophrenia is proposed to result from the abnormal hierarchical cerebral organization rather than from individual systems alone (Yang et al., 2016). However, detecting abnormality in the cerebral hierarchy organization represents a challenge, due to the limited number of approaches explicitly designed to evaluate hierarchical information propagation in the brain system.

Recent advances in neuroscience towards the understanding of brain organizational principles have highlighted a cortical hierarchy as a unifying functional mechanism for information processing in the primate and mouse brains (Burt et al., 2018; Chaudhuri et al., 2015; Demirtaş et al., 2019; Fulcher et al., 2019; Margulies et al., 2016; Paquola et al., 2020, 2019; Taylor et al., 2015). Specifically, this mechanism refers to the functional system extending from primary sensorimotor to association areas, along which it increasingly represents more abstract and complex information in the brain (Fulcher et al., 2019; Huntenburg et al., 2018; Margulies et al., 2016; Mesulam, 2012; Taylor et al., 2015). This hierarchical architecture facilitates segregated processing of specialized function domains (e.g., sensory and cognitive process), while also enabling a dynamic configuration and cross-communication of networks for more complex and integrated mental activity (Huntenburg et al., 2018; Murphy et al., 2018; Sepulcre et al., 2012; Taylor et al., 2015). Investigating the cortical hierarchy provides an integrative window into the impairments of functional integration between and within early sensory processing and high order cognitive functions in schizophrenia.

The present study aimed to examine how the impaired function and integration of both sensory and cognitive processes relates to the macroscale cortical hierarchy in schizophrenia. We applied a novel combination of connectome gradient mapping (Margulies et al., 2016) and stepwise functional connectivity (SFC) analyses (Sepulcre et al., 2012), which offer a complementary characterization of hierarchical abnormalities in schizophrenia. In other neuropsychiatric conditions, such as autism spectrum disorder, such hierarchy sensitive techniques have recently been able to identify connectome-level substrates that can similarly account for remarkable heterogeneity at the phenotypic level (Hong et al., 2019). Specifically, gradient mapping approach allows the visualization of spatial trends in connectivity variations following the cortical hierarchy. This approach describes a continuous coordinate system at the systems level that place sensory and motor networks on one end and transmodal network on the other. This approach thus provides us a simplified representation in terms of main dimensions to characterize the alteration of the macroscale cortical hierarchy in schizophrenia. SFC, developed earlier, has shown that cortical hierarchy can also be understood as a sequence of steps in connectivity space. SFC was initiated from a priori-defined sensory seeds-based FC to further examine the hierarchical stream of information from unimodal sensory regions (visual, auditory, and somatosensory) to transmodal regions in schizophrenia patients and healthy controls (HC). This approach thus enabled us to investigate the presence of atypical functional transitions from unimodal to multimodal cortical areas within the framework of cortical hierarchy in schizophrenia. Therefore, SFC is well-suitable to understand the ‘bottom-up’ dysregulation of higher cognitive functions in this condition mentioned above.

Based on recent models emphasizing the ‘bottom-up’ dysregulation of higher cognitive functions (Javitt, 2009a; Javitt and Freedman, 2015; Javitt and Sweet, 2015) and the initial observation of impaired communication between sensory and cognitive processes system in schizophrenia (Berman et al., 2017; Hoptman et al., 2018; Kaufmann et al., 2015), we hypothesized that the pathological interaction between sensory and cognitive processes in schizophrenia would be reflected by the expected observation of the prominent alteration of somatosensory-motor system in the hierarchical stream of information flows propagated in the brain of individuals with schizophrenia. Also, we hypothesized that the pathological interaction between sensory and cognitive processes in schizophrenia would be reflected by the expected abnormal communication between the sensorimotor and high order cognitive system in the hierarchical organization of the brain in individuals with schizophrenia.

## 2. Methods

### 2.1 Participants

One hundred and two patients with schizophrenia were recruited from the Clinical Hospital of Chengdu Brain Science Institute; 126 HC were recruited from the local community through advertisements and word of mouth. Patients were diagnosed with schizophrenia according to the structured clinical interview for DSM-IV Axis I disorders - clinical version (SCID-I-CV). All patients received treatment with antipsychotics and did not participate in other therapy programs. Exclusion criteria included co-morbid Axis I diagnosis, active substance use disorders, or history of brain injury. HC were excluded based on current or past Axis I disorder as verified using the Structured Clinical Interview for DSM-IV, history of neurological illness, traumatic brain injury, substance-related disorders, or first-degree relatives with a history of psychosis. Two HC with poor quality of imaging data as assessed by visual evaluation were excluded. Six patients and two HC were further excluded based on the result of MRI preprocessing (see the method for details). This process left ninety-six schizophrenia patients and 122 HC as a final sample of our study. Written informed consent was obtained from all healthy participants and the patients or guardians of patients. This study was approved by the Ethics Committee of the Clinical Hospital of Chengdu Brain Science Institute following the Helsinki Declaration of 1975, as revised in 1978. Demographic and clinical information of all participants is shown in Table 1. Two groups did not show statistically significant differences in age, sex, education and handedness (p>0.3). The mean illness duration is 15.1 years.

**Table 1.**
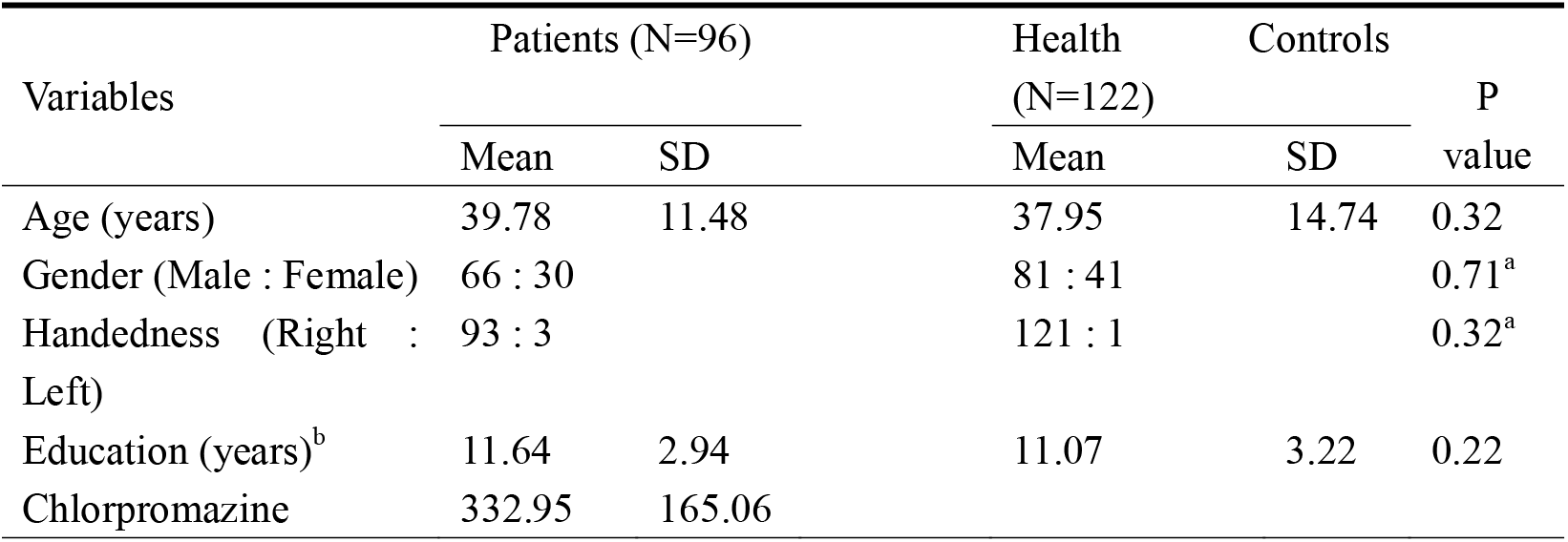
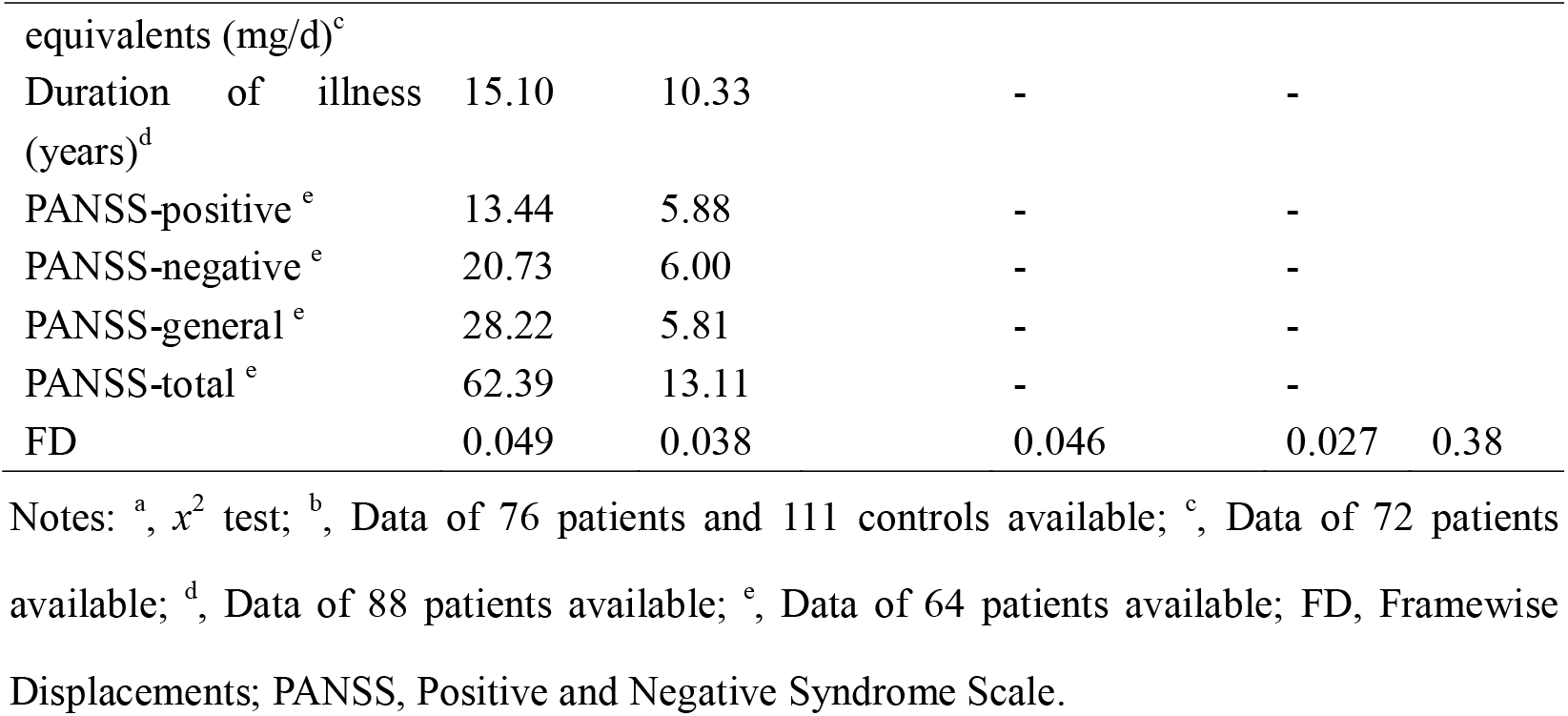
Demographic characteristics of schizophrenia patients and controls.

### 2.2 Data acquisition and image preprocessing

MRI data were obtained on a 3-T GE Discovery MR 750 scanner at the MRI Center of the University of Electronic Science and Technology of China. Functional scans were performed using a standard echo-planar imaging (EPI) pulse sequence with the following scan parameters: TR/TE = 2000 ms/30 ms, flip angle (FA) = 90°, matrix size = 64 × 64, field of view (FOV) = 240 × 240 mm^2^, 35 interleaved slices and slice thickness = 4 mm (no gap). During this resting-state fMRI scanning, each participant was instructed to stay relaxed and close his/her eyes without falling asleep. Each scan lasted for 510 s per subject (255 volumes). T1-weighted anatomical data were acquired using a MPRAGE (MEMPR) sequence (scan parameters: TR/ TE= 1900 ms/3.43 ms, FA = 90°, matrix size = 256 × 256, FOV = 240 mm × 240 mm, slice thickness = 1 mm, voxel size = 0.9375 mm × 0.9375 mm × 1 mm, 160 slices). In both scans, foam pads were used to reduce head movement and scanner noise. The anatomical data were used to normalize functional data.

All preprocessing steps were carried out using the Data Processing & Analysis for (Resting-State) Brain Imaging (DPABI v4.1, (Yan et al., 2016)) and Matlab scripts. Consistent with our previous study (Dong et al., 2018a), functional images were (1) discarded in the first five volumes, (2) slice-time corrected, (3) realigned, (4) co-registered to the high-resolution 3D anatomic volume, (5) warped into MNI152 standard space (resampling the voxel size into 3×3×3 mm^3^), (6) underwent wavelet despiking of head motion artifacts (Patel et al., 2014)), (7) underwent regression of motion and non-relevant signals, including linear trend, Friston 24 head motion parameters (Friston et al., 1996; Satterthwaite et al., 2013), white matter (CompCor, 5 principal components), and CSF signal (CompCor, 5 principal components, (Behzadi et al., 2007)), and (8) were filtered using a band-pass filter (0.01-0.1 Hz). We excluded participants due to maximum head motion exceeding 2.5 mm or 2.5° rotation or with >10% frame-to-frame motion quantified by framewise displacements (FD>0.5mm, (Power et al., 2012))) during MRI acquisition. Besides, mean FD was evaluated in the two groups (Power et al. 2012). The mean FD for each participant was evaluated using the following formula:

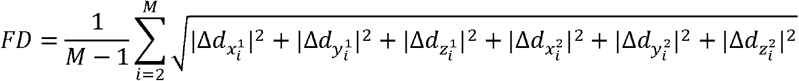

where M is the number of the fMRI time points, and 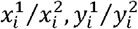 and 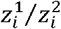 are translations/rotations at the *i*th time point in the *x,y* and *z* directions, respectively; 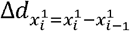. The global mean signal was not regressed out because this may distort between-group comparisons of inter-regional correlation (Saad et al., 2012). Besides, studies suggest that altered global signal is an important neuroimaging feature in schizophrenia (Hahamy et al., 2014; Yang et al., 2014). To reduce computational demands, the rsfMRI data were down-sampled to 4 mm isotropic voxels, resulting in 18815 voxels. Subsequent gradient and SFC analyses were voxel-based calculation. Consistent with previous voxelwise FC computation (Tomasi and Volkow, 2010), spatial smoothing was not conduct in the image preprocess step, but was conducted in postprocessing steps (6 mm) for gradient and SFC maps.

### 2.3 Connectivity gradient analyses

Gradient mapping techniques describe a continuous coordinate system at the systems level that place sensory and motor networks on one end and transmodel network on the other. This approach thus provides us a simplified representation in terms of main dimensions to characterize the alteration of the macroscale cortical hierarchy in schizophrenia.

Specificlly, the voxel-level connectivity matrix for each subject was computed using Fisher Z-transformed Pearson correlations. Based on previous studies (Dong et al., 2020; Guell et al., 2018; Hong et al., 2019; Margulies et al., 2016; Vos De Wael et al., 2018), we thresholded the rsFC matrix with the top 10% of connections per row retained, whereas all others were zeroed. The negative connections were zeroed as well. Then, we used cosine distance to generate a similarity matrix that reflected the similarity of connectivity profiles between each pair of voxels. We used diffusion map embedding (Coifman et al., 2005), a nonlinear dimensionality reduction algorithm, to identify a low-dimensional embedding from a high-dimensional connectivity matrix. This methodological strategy has been proved to successfully identify relevant aspects of functional organization in the cerebral cortex in previous studies (Hong et al., 2019; Margulies et al., 2016). Similar to Principal Component Analysis (PCA), diffusion map embedding can identify principal gradient components accounting for the variance in descending order. The result of diffusion embedding is not one single mosaic of discrete networks, but multiple, continuous maps (gradients), which capture the similarity of each voxel’s functional connections along with a continuous space. All gradients are orthogonal to each other and capture a portion of data variability in descending order.

There is a single parameter αto control how the density of sampling points affects the underlying manifold (α = 0, the maximal influence of sampling density; α = 1, no influence of sampling density) in the diffusion map embedding algorithm. Following previous studies (Guell et al., 2018; Hong et al., 2019; Margulies et al., 2016), we set α = 0.5, which can help retain the global relations between data points in the embedded space. To compare between the SZ and HC groups, we used an average connectivity matrix calculated from all patients and controls to produce a group-level gradient component template. We then performed Procrustes rotation to align the gradients of each participant to this template, following the strategy of previous analyses (Langs et al., 2015). To maximize interpretability, we only used the first gradient component in our analyses. The first gradient explains as much of the variance in the data as possible (Figure S1) and, from a neurobiological point of view, represents a well-understood sensorimotor-to-transmodal organizational principle in the cerebral cortex connections.

### 2.4 Stepwise functional connectivity analyses

SFC analysis is a graph-theory-based method that detects both direct and indirect functional couplings from a defined seed region to other regions in the brain. More importantly, SFC analytical approach allows for analysis of indirect FC (medium and large connectivity distances from the seed), which is thought to provide information integration about hierarchical flow across specific brain networks (Sepulcre, 2014; Sepulcre et al., 2012). This approach thus enabled us to investigate the presence of atypical functional transitions from unimodal to multimodal cortical areas within the framework of the cortical hierarchy in schizophrenia.

SFC analysis computes the number of functional paths between defined seed regions and every other voxel in the brain at successive numbers of relay stations or “link-step” distances (Sepulcre, 2014; Sepulcre et al., 2012). Hence, it complements connectivity gradient approaches by allowing voxel-level functional connections to be assessed at a range of intermediate relay stations. Following previous studies (Martínez et al., 2019; Sepulcre et al., 2012), connectivity matrices were first filtered to include only positive correlations due to the ambiguous interpretation of negative correlations. After that, the connectivity matrices were further filtered to contain only correlations surviving a stringent false discovery rate (FDR) correction (q < 0.001). Finally, we submitted the resulting FDR thresholded matrices to SFC analysis.

Given that deficits of visual, auditory, and somatosensory processing in schizophrenia were consistently observed (Javitt, 2009b; Javitt & Freedman, 2015, for reviews), three bilateral primary sensory seed regions of interest (ROIs) including visual (MNI coordinate x, y, z: −14/10 [left/right], −78, 8; (Brodmann 17, V1)), auditory (−54/58, −14, 8; (Brodmann 22, A1)) and somatosensory (−42/38, −29, 65; (Brodmann 3, hand area)) areas (Sepulcre et al., 2012), were defined as cubic regions of eight voxels each. To assess the degree of combined SFC of all sensory seeds irrespective of modality, a combined mask was constructed by combining information from all three primary sensory regions. The method is described in detail elsewhere (Sepulcre, 2014; Sepulcre et al., 2012) and schematically represented in Figure 1.

**Figure 1.**
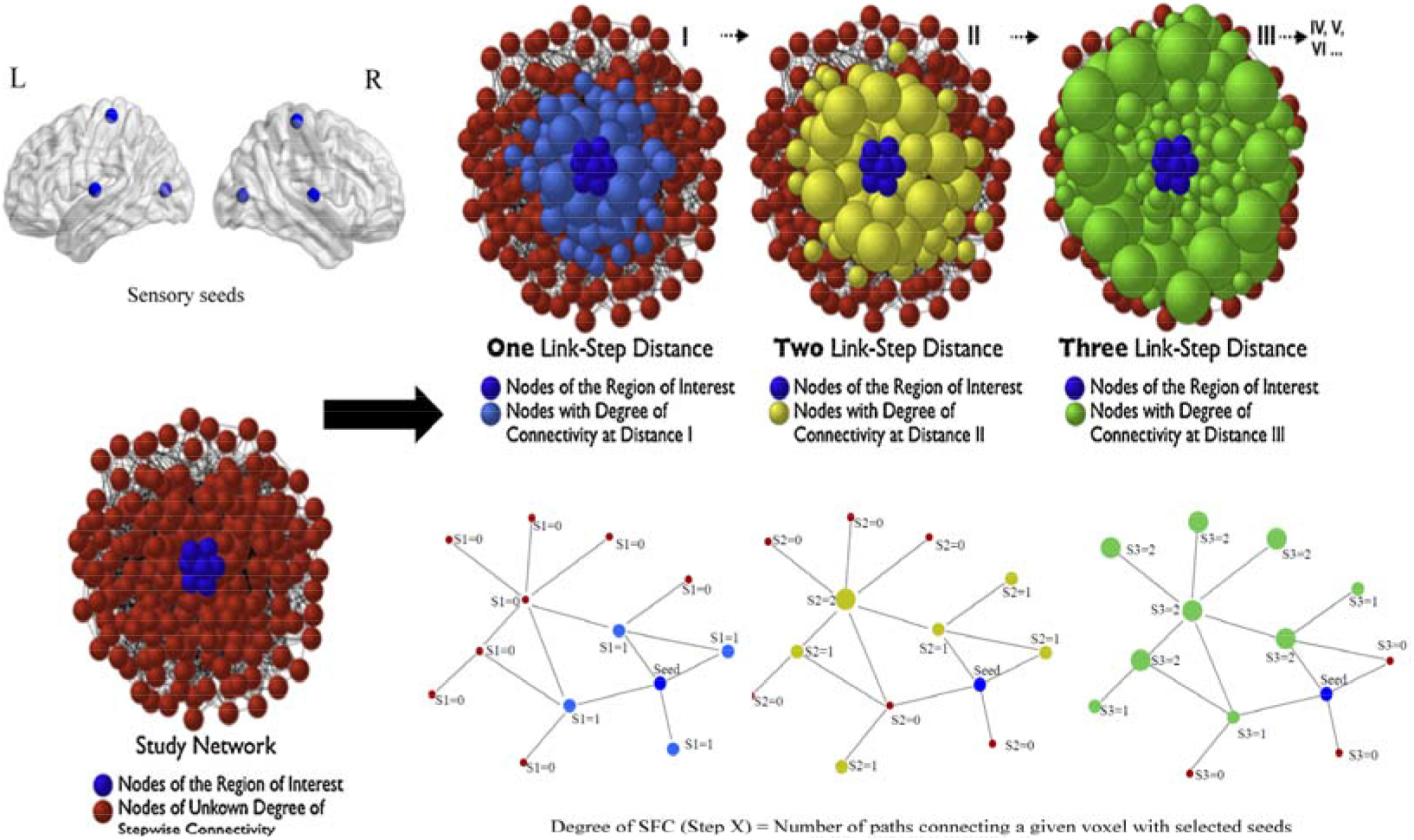
Computation of the SFC degree. Figure adapted from Sepulcre (2014) describing the SFC analysis. (Reproduced with permission from Elsevier, Integration of visual and motor functional streams in the human brain, Neuroscience Letters (2014), 567: 68-73.)

The degree of SFC of a given voxel of the brain is defined as the number of functional paths connecting that voxel with an a priori selected seed region at a specific link-step distance. A link-step distance is defined as the number of edges that pertain to a path connecting a given voxel to the seed regions. At each link step, SFC maps were standardized to Z-scores by subtracting the mean and dividing by its standard deviation (SD) to yield SFC values. Therefore, each SFC map represents a relative increase of connectivity degree across different link-step distances. As demonstrated in previous studies (Buckner et al., 2009; Sepulcre et al., 2012), functional pathways “collapse” into the cortical hubs of the adult human brain after link-step distances >7; accordingly, we constrained our SFC analysis to seven link-step distances.

### 2.5 Statistical and Control Analyses

To visualize the gradient pattern, group mean maps were calculated for each group. One-sample t-tests were performed to characterize the SFC patterns at each of the seven link-step distances in the HC and schizophrenia groups separately (p<0.001 uncorrected, only for purposes of clear data visualization).

Two-sample t-tests were calculated to determine diagnostic differences (schizophrenia patients (SZ) Vs. HC) in Z-normalized values of the principle (first) gradient scores, and SFC degree at each of the seven link-step distances. Age, sex, handedness and mean FD were set as covariates. Results for each test are reported at a voxel-based threshold corrected for false discovery rate of multiple comparisons (FDR voxel-wise correction p< 0.05). We also imposed a minimum cluster extent of 20 voxels. All the results reported below are based on without global signal regression.

Three analyses were performed to ensure robustness of the main findings. First, because GSR is controversial, we repeated core analyses (gradient and SFC) with GSR. Second, as shown in Table 1, while there was no significant difference in mean framewise displacement (FD) between patients and controls, we also corrected for head motion in the subsequent statistical comparisons by using mean FD as covariate (Yan et al., 2013). And, to investigate the potential effects of micro head motion on our findings, we calculated Pearson Correlations between altered gradients, SFC value and mean FD. Third, to target the potentially confounding effect of medication, we calculated Pearson Correlations between altered gradients, SFC value and medication measured by chlorpromazine equivalents.

### 2.6 Correlations between altered gradient scores, SFC degree, and clinical variables

To further examine the relationship between altered gradient scores, SFC degree, and clinical features, we calculated Pearson Correlations between gradient scores, SFC degree and the severity of clinical symptoms measured by PANSS (positive, negative, general psychopathology symptoms subscales and overall scores) in the patients’ group. Analyses were computed in each region where the SZ and HC groups differed significantly in the two-sample *t*-tests. Given its high correlation with age, illness duration was not included separately in correlation analysis (Moser et al., 2018).

### 2.7 Data and code availability

The preprocessing software is freely available (DPABI v4.1, http://rfmri.org/dpabi). The code for gradient analysis is openly available via the BrainSpace toolbox (http://brainspace.readthedocs.io) (Wael et al., 2020). The code for SFC analysis is available via a direct request to Jorge Sepulcre. Results were visualized with BrainNet Viewer v1.7 (https://www.nitrc.org/projects/bnv/) (Xia et al., 2013). The imaging and clinical data are made available via a direct request to the corresponding author (Cheng Luo). Sharing and re-use of imaging and clinical data need the expressed written permission of the authors and clearance from the relevant institutional review boards.

## 3. Results

### 3.1 The principal functional gradient of cerebral cortex in schizophrenia

The principal gradient of cerebral cortex FC showed a similar sensorimotor-to-transmodal gradient of cortical organization in HC and SZ (Figure 2). It extended from primary cortices to transmodal areas. Of note, there was no significant difference between SZ and HC in the explained variance of the principle gradient (two-sample t-test, *t* = 0.86, *p* = 0.39).

**Figure 2.**
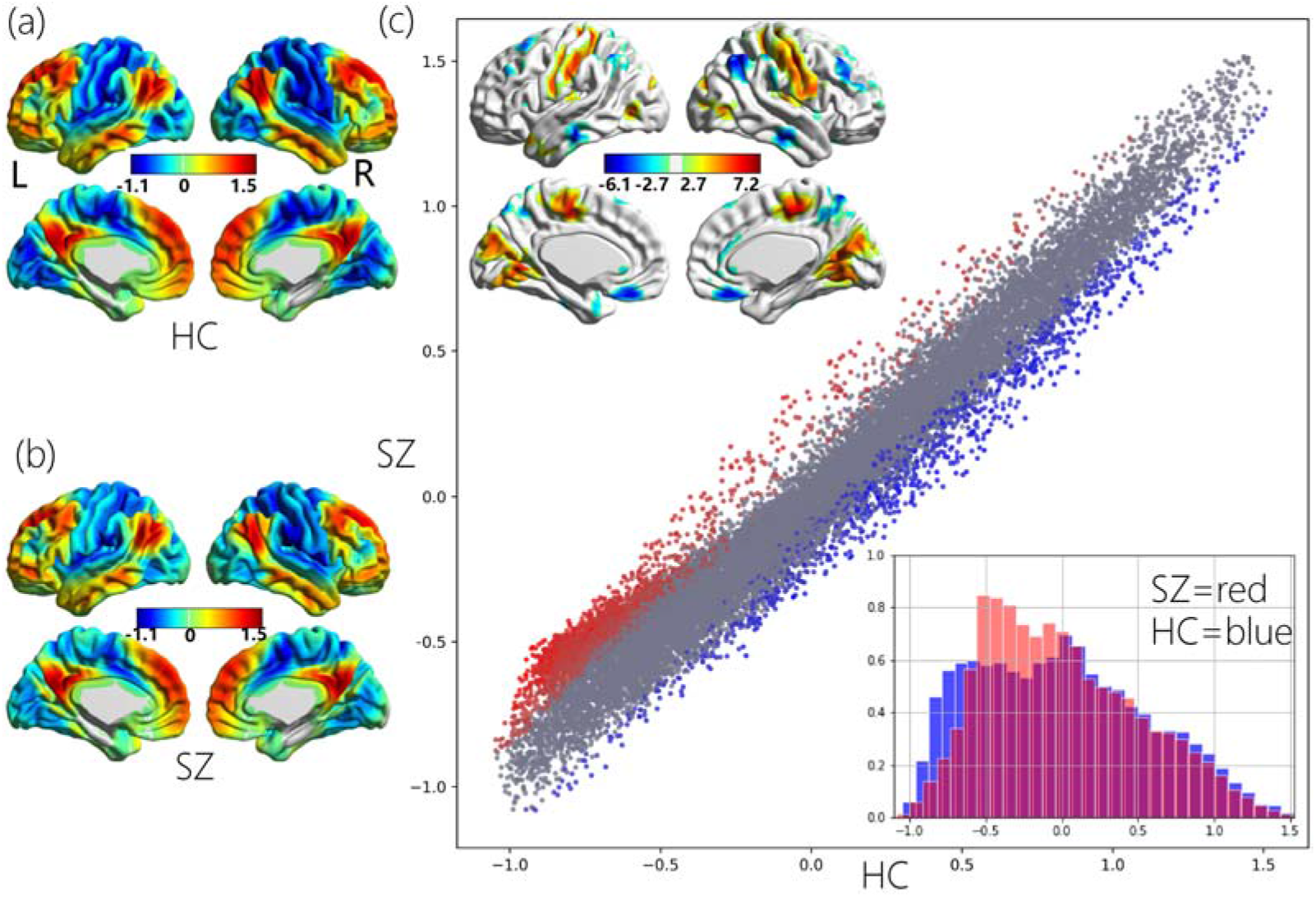
Group patterns and differences in the cerebral principal functional gradient. (a) Gradient pattern in HC. (b) Gradient pattern in SZ. (c) Group differences between SZ and HC. Scatterplot represents cerebral gradient of SZ (y axis) vs. cerebral gradient of HC (x axis).

Scatterplot colors correspond to group differences map as shown in top left corner of Figure 2(c): higher gradient value in SZ (red), and lower gradient value in SZ (blue) compared to HC. Compressed gradient pattern in SZ is shown in density histograms in bottom right corner of Figure 2(c). All results are shown after FDR correction (P < 0.05).

Compared to HC (Figure 2 and Table 2), Schizophrenia patients showed increased gradient values in regions of sensorimotor network and visual network, including bilateral post/precentral gyrus, posterior insula, middle occipital gyrus, and lingual gyrus; and decreased gradient scores in transmodal regions mainly belonging to FPN, i.e., middle frontal gyrus, superior frontal gyrus, inferior parietal lobule; also including a few regions in DMN (e.g., medial frontal gyrus, middle temporal gyrus and angular gyrus). As shown in the scatterplot of Figure 2c, functional gradient abnormalities in this case extended across the whole principle gradient spectrum. More specifically, higher principle gradient values in the SZ group were localized in the lowest pole of principle gradient (which corresponds to primary sensorimotor and visual processing areas), whereas lower values in the SZ group extended from the medium aspects to the highest pole of principle gradient (transmodal regions).

**Table 2.**
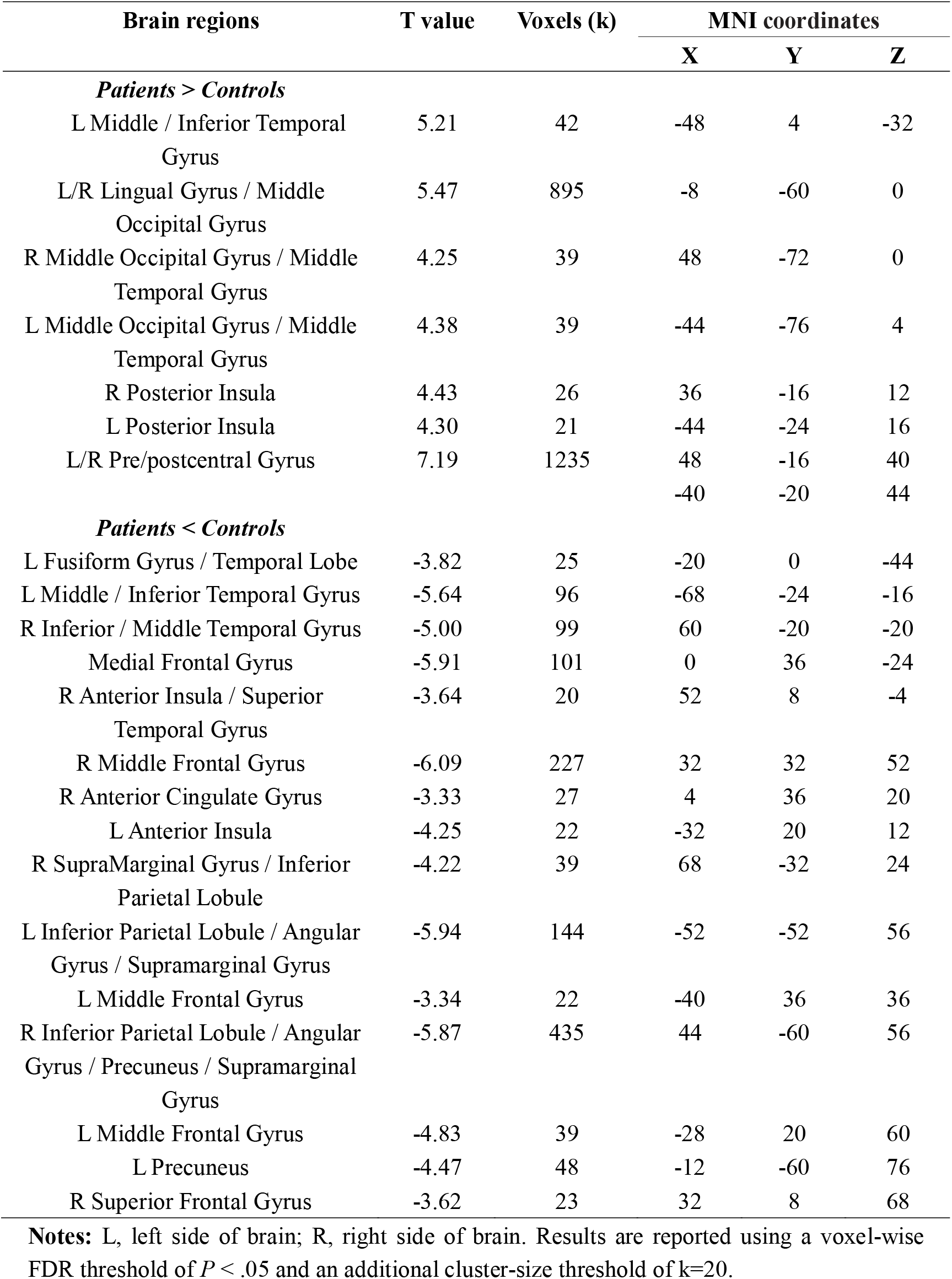
Group differences in principal functional gradient values.

| Brain regions | T value | Voxels (k) | MNI coordinates |  |  |
| --- | --- | --- | --- | --- | --- |
|  |  |  | X | Y | Z |
| *Patients > Controls* |  |  |  |  |  |
| L Middle / Inferior Temporal Gyrus | 5.21 | 42 | -48 | 4 | -32 |
| L/R Lingual Gyrus / Middle Occipital Gyrus | 5.47 | 895 | -8 | -60 | 0 |
| R Middle Occipital Gyrus / Middle Temporal Gyrus | 4.25 | 39 | 48 | -72 | 0 |
| L Middle Occipital Gyrus / Middle Temporal Gyrus | 4.38 | 39 | -44 | -76 | 4 |
| R Posterior Insula | 4.43 | 26 | 36 | -16 | 12 |
| L Posterior Insula | 4.30 | 21 | -44 | -24 | 16 |
| L/R Pre/postcentral Gyrus | 7.19 | 1235 | 48 | -16 | 40 |
|  |  |  | -40 | -20 | 44 |
| ***Patients < Controls*** |  |  |  |  |  |
| L Fusiform Gyrus / Temporal Lobe | -3.82 | 25 | -20 | 0 | -44 |
| L Middle / Inferior Temporal Gyrus | -5.64 | 96 | -68 | -24 | -16 |
| R Inferior / Middle Temporal Gyrus | -5.00 | 99 | 60 | -20 | -20 |
| Medial Frontal Gyrus | -5.91 | 101 | 0 | 36 | -24 |
| R Anterior Insula / Superior Temporal Gyrus | -3.64 | 20 | 52 | 8 | -4 |
| R Middle Frontal Gyrus | -6.09 | 227 | 32 | 32 | 52 |
| R Anterior Cingulate Gyrus | -3.33 | 27 | 4 | 36 | 20 |
| L Anterior Insula | -4.25 | 22 | -32 | 20 | 12 |
| R SupraMarginal Gyrus / Inferior Parietal Lobule | -4.22 | 39 | 68 | -32 | 24 |
| L Inferior Parietal Lobule / Angular Gyrus / Supramarginal Gyrus | -5.94 | 144 | -52 | -52 | 56 |
| L Middle Frontal Gyrus | -3.34 | 22 | -40 | 36 | 36 |
| R Inferior Parietal Lobule / Angular Gyrus / Precuneus / Supramarginal Gyrus | -5.87 | 435 | 44 | -60 | 56 |
| L Middle Frontal Gyrus | -4.83 | 39 | -28 | 20 | 60 |
| L Precuneus | -4.47 | 48 | -12 | -60 | 76 |
| R Superior Frontal Gyrus | -3.62 | 23 | 32 | 8 | 68 |

**Notes:** L, left side of brain; R, right side of brain. Results are reported using a voxel-wise FDR threshold of $P < .05$ and an additional cluster-size threshold of $k=20$ .

To better characterize the altered pattern of sensorimotor-transmodal hierarchical gradient, global histogram analyses were performed. As shown in the right bottom corner of Figure 2c, this analysis revealed that there was a prominent compression of the lowest portion of the principal functional gradient and a less prominent compression of the highest portion of the principal functional gradient. Further, Kolmogorov-Smirnov test (Matlab function) indicated that the distribution of SZ group gradient values was significantly different from the distribution of HC group gradient values (p<0.001, ks2stat=0.08). To quantitatively demonstrate the overall compression, we tested whether there was a linear correlation between X and X-Y per corresponding voxel (X=SZ group gradient values, i.e., red histogram of Figure 2c, Y=HC group gradient values, i.e., blue histogram of Figure 2c). We found there was a significant correlation between X-Y and X (r=0.51, p<0.001), which suggested the overall gradient value compression in SZ group compared to HC group. In the same logic, we tested the compression of transmodal pole and sensorimotor pole, we found the correlation value in sensorimotor pole (r=0.53, p<0.001) was higher than transmodal pole (r=0.10, p<0.001), which suggested there was a prominent compression of sensorimotor regions and a less prominent compression of transmodal regions.

Connectivity gradient analysis provides a description of the connectome where each voxel is located along a gradient according to its connectivity pattern. Voxels with similar connectivity patterns are located close to one another along a given connectivity gradient. Therefore, the gradient value represents information about the spatial pattern in the embedding space - shifts in value are not ‘more’ or ‘less,’ but rather reflect changes in relative similarity within a latent dimension, i.e., the similarity of functional connectivity patterns along each dimension (‘gradient’). The gradient values are a scalar, and for this reason significant gradient value alterations in schizophrenia reflect the extent to which the patient group deviates from the HC group. Our interest here was the different spatial distributions of cortical hierarchy between two groups. Interestingly, our finding of compressed cerebral cortical functional gradients suggested a less differentiated global hierarchical organization, i.e., diminished network differentiation in schizophrenia, in which there is a relatively stronger shift in functional affiliation from visual-sensorimotor towards transmodal regions in gradient space.

### 3.2 The SFC degree in schizophrenia

The SFC degree showed a similar spatial transition pattern along the sensation-to-cognition continuum in SZ and HC (Figure 3a). In steps 1 and 2, sensory-related seeds display a regional-local FC along with the unimodal areas, e.g., somatomotor and visual cortex. From 3 to 7 link-step distances, sensory-related seeds showed gradual transitions towards multimodal integration areas (e.g., dorsal anterior cingulate cortex and frontal eye field, frontoinsular cortex), and eventually displayed convergent to the cortical hubs regions (e.g., dorsolateral prefrontal, inferior lateral parietal cortex, medial prefrontal cortex, posterior cingulate cortex / precuneus or the lateral temporal cortex).

**Figure 3.**
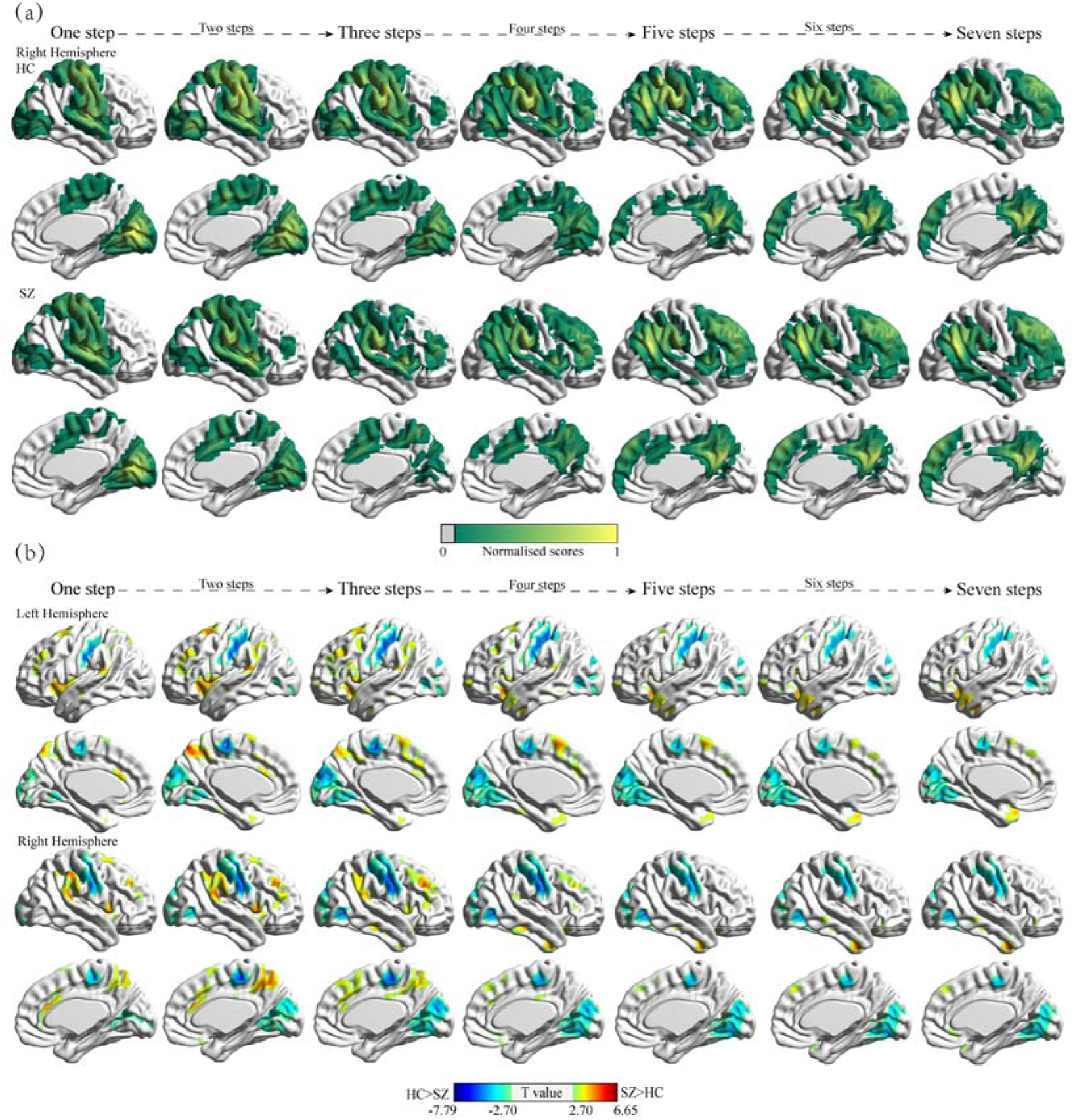
Group patterns and differences in stepwise functional connectivity degree. (a) Stepwise functional connectivity patterns in HC and patients with schizophrenia (One-Sample t Tests with p<0.001 uncorrected). In the normalized color scale, 0 represents nonsignificant results (p < 0.001), and 1 is the maximum value corresponding to the smallest p value. Given the results of the left hemisphere are similar to those on the right hemisphere, we only show the results of the right hemisphere for visualization (b) Group differences between schizophrenia and HC in stepwise functional connectivity degree. All results are shown after FDR correction (P < 0.05).

Statistical comparison indicated that at link-step distance 1, corresponding to classic seed-based FC analysis, patients with schizophrenia showed reduced SFC degree between the unimodal seeds and visual and sensorimotor systems, i.e., bilateral middle occipital gyrus, lingual gyrus and pre/postcentral gyrus (Table S1 and Figure 3b). Interestingly, this reduced SFC pattern was consistently observed across all link-steps distances (Step 2-7). Increased SFC degree was found between unimodal seeds and frontoparietal regions, i.e., middle / superior frontal gyrus, inferior parietal lobule, supramarginal gyrus, and dorsal precuneus), and ventral attention regions (dorsal anterior cingulate cortex and bilateral anterior insular cortex / central opercular cortex) at early and medium link-step distances (Step1 to 4). However, this increased SFC degree gradually faded in the remaining link-step maps (steps 5 to 7).

### 3.3 Control analyses

We summarize three analyses that ensured robustness of results. GSR did not significantly affect trends of overall results (gradient and SFC analyses). Detained discussion can be found in Supplementary Material (Figure S2-3). The relatively consistent results between without GSR and with GSR indicated the observed main findings reflected the reliable pathophysiology of schizophrenia. Second, we found that FD was not associated with altered gradient and SFC degree scores (all p values in this analysis were larger than 0.05), indicating that group differences reported here are rather unlikely to be driven by head motion. Similarly, Chlorpromazine equivalents were not associated with an altered gradient and SFC degree (all p >0.1), suggesting that these changes are unlikely to be mainly driven by medication.

### 3.4 Association between altered gradient, SFC and clinical severity of symptoms in schizophrenia

The severity of clinical symptoms was related to decreased functional gradient values in ventral medial frontal gyrus, left anterior insula and left precuneus (Table S2). Across all the link-step distances, most of the significant correlations fit the following rule: for those regions involved in high order cognitive function (i.e., superior frontal gyrus, anterior insular cortex), increased SFC degree was associated with less clinical severity. Accordingly, for those regions involved in sensory processing function, i.e., pre/postcentral gyrus and visual areas, increased SFC degree correlated with greater clinical severity (uncorrected). There is an exception in the left middle frontal gyrus at step 4, where increased SFC degree was associated with greater clinical severity.

## Discussion

Recent emerging models and rapidly growing empirical studies emphasize the impaired function and integration of both the early sensory and cognitive processing in understanding the pathophysiology of schizophrenia. Investigating the fundamental sensorimotor-to-transmodal cortical hierarchy organization in schizophrenia would provide critical and integrative experimental evidence for these models. The present study used a novel combination of connectome gradient and SFC analyses to characterize the macroscale cortical hierarchy organization in schizophrenia. In summary, the gradient analysis identified a significantly reduced network differentiation, i.e., gradient compression, in which there is a prominent compression from the sensorimotor system of the cortical hierarchy and a less prominent compression from the higher-level systems such as FPN and DMN. The SFC approach further suggested reduced network differentiation related to atypical functional transitions from unimodal to multimodal cortical areas in schizophrenia. Altogether, the present study provided converging evidence for abnormal cortical hierarchy organization as a system-level substrate underlying the pathological interaction of early sensory and cognitive function in schizophrenia. The findings indicated that impairments at different hierarchical processing steps, especially the foci of effects emphasize that disrupted somatosensory-motor systems may cascade into the higher-order cognitive deficits, which are the hallmark characteristic of schizophrenia.

### 4.1 The cascading effects of early sensory deficits along the sensation-to-cognition continuum

Intriguingly, the present study found a selective gradient compression pattern of the sensorimotor system in the compressed cortical hierarchy organization. Recently, a significant paradigm shift in research filed of schizophrenia has begun to emerge, according to which early regions of the sensory pathway may result in ‘bottom-up’ dysregulation of higher cortical function (Javitt, 2009a, 2009b; Javitt and Freedman, 2015; Javitt and Sweet, 2015). Further complementing this view, neurophysiology findings indicate that the bottom-up propagation of deficits from early sensory to higher-level processes in schizophrenia occurs even when top-down processes remain intact (Dias et al., 2011). Similarly, some recent rsFC and structural studies found that the connectivity deficits of visual and sensorimotor pathway can be detected even when the associative regions, like FPN and DMN, failed to reach statistical significance in schizophrenia (Bordier et al., 2018; Chen et al., 2014; Guo et al., 2014; Jørgensen et al., 2016; Liu et al., 2018; Youxue Zhang et al., 2019). Our findings of prominent cortical network compression in the sensorimotor and visual systems therefore provides an integrative basis to support previous reports of impaired early sensory processing in schizophrenia. More importantly, this result extends previous findings by showing that the abnormality of early sensory processing is not purely anchored on local functional circuits but rather affects overarching hierarchical cerebral organization, which reflects its global and widespread influence on brain information transitions in schizophrenia.

Furthermore, the SFC findings strengthened global and widespread influence of the early sensory processing on brain information transitions in schizophrenia by demonstrating that the pattern of hypo-connectivity within visual and sensorimotor systems across all (early, medium and large) link-step distances along the sensation-to-cognition continuum. In line with the prominent compression from the sensorimotor and visual system, our stable observation of hypo-connectivity within visual and sensorimotor systems across all link distances suggests an impaired ability to decode early information coming from early sensory regions in schizophrenia. Moreover, the degraded quality of early sensory input would persist, propagate and form the cascading effects along the whole sensation-to-cognition continuum, affecting higher-order integrative networks at subsequent stage of the cortical hierarchy in schizophrenia. These findings provide novel and integrative evidence to support the propagation of sensory deficits to higher cognitive functions in schizophrenia (Calderone et al., 2013; Dondé et al., 2019b; Javitt, 2009a; Leitman et al., 2010).

In addition, although deficit of early sensory processing systems is not emphasized within prevailing psychiatric models, our findings are in parallel to the recent neurobiological findings of abnormal early sensory processing which characterize individuals’ variability in psychopathology and cognitive impairment across multiple psychiatric disorders (Elliott et al., 2018; Kebets et al., 2019). So far, with some notable exceptions in neurophysiological studies, hypothesis-driven fMRI studies have primarily targeted high-order brain networks. Consistent with the previous observation of prominent motor pole compression in cerebellar hierarchy (Dong et al., 2020), the present data-driven findings encourage future research to expand the neuroscientific view in schizophrenia and give more attention to sensory processing deficits characterization. Further investigation along these lines would deepen our understanding of the pathophysiology of schizophrenia.

### 4.2 Inefficient integration of bottom-up sensory information with top-down processes

We also found that schizophrenia patients showed hyper-connectivity between unimodal sensory seeds and frontoparietal-ventral-attention regions at the early and medium link distances along the sensation-to-cognition continuum. Several previous seed-based or ROI-based FC studies provide preliminary evidence for sensory networks pathologically interaction with higher-order association systems in the literature of schizophrenia (Berman et al., 2017; Hoptman et al., 2018; Kaufmann et al., 2015). In a more holistic view, we extend the previous studies by showing that this pathological interaction between sensory networks and higher-order association systems exists in the hierarchical information flow, not limited in directed communication (1 link-step). Supporting the cascading effects of sensory process deficits on subsequent high-order cognitive process impairment, our hyper-connectivity may further suggest that the integration of bottom-up sensory information with attentional demands and executive control processes in the sensation-to-cognition continuum is more effortful or less efficient in schizophrenia than in healthy populations.

This inefficient integration process is also characterized by the overall compression of the principal sensorimotor-to-transmodal hierarchy organization, which reflects diminished separation between sensory systems (e.g., visual and sensory regions) involved in the immediate environment and transmodal cognitive systems (e.g. frontoparietal regions) that support complex cognitive inferences. The effective brain function is supported by the maintenance of subnetworks segregation as well as their integration (Wig, 2017). Therefore, the diminished network differentiation would unavoidably result in ineffective functional specialization, leading to a blurred boundary between externally oriented immediate environment and internally abstract cognitive processing (Murphy et al., 2018; Northoff and Duncan, 2016), which further contributes to the inefficient integration of bottom-up sensory information with top-down processes.

### 4.3 An integrative/holistic neuroscientific perspective to understand the ‘bottom-up’ dysregulation of higher cognitive function

It is recent that neurobiological accounts of schizophrenia have begun to put underline on changes in aberrant sensory (bottom-up) processing, although this deficit has already been suggested earlier. It has been evidenced that the sensory and perceptual deficits can be attributed as the significant role in higher-level cognitive processes and clinical symptoms (Javitt and Freedman, 2015). So far, hypothesis-driven fMRI studies have primarily targeted high-order cognitive circuits, such as frontal-parietal and attention regions as the neural correlates of high-order cognitive deficits (Barch and Ceaser, 2012). Although previous neurophysiological studies conclude developmental effects of the impaired sensory process on high-order functions through a gradual (some even termed of ‘hierarchical’) propagation way in schizophrenia (Butler et al., 2007; Calderone et al., 2013; Dias et al., 2011; Dondé et al., 2019b; Hoptman et al., 2018; Leitman et al., 2010), the potential neural correlates of the hierarchical propagation of sensory deficits to higher cognitive functions remained elusive. Using a novel combination of connectome gradient and SFC analyses, investigating the fundamental sensorimotor-to-transmodal cortical hierarchy organization in schizophrenia provided critical and integrative neural correlates of pathological interaction of early sensory and cognitive function, especially ‘bottom-up’ dysregulation of higher cognitive function in schizophrenia. That is, within the framework of compressed cortical hierarchy organization, a cascade of impairments stemming from the disrupted somatosensory-motor system and inefficient integration of bottom-up sensory information with attentional demands and executive control processes may partly account for high-level cognitive deficits of schizophrenia. Critically, these abnormalities showed trend associations with the severity of clinical symptoms, further highlighting the importance of cascading impairments of sensory processing and less efficient integration between sensory and cognitive processing to understand the clinical profiles of schizophrenia. Considering most of these associations cannot survive after multiple comparison correction, it should keep cautious to interpret these trend symptom associations. While top-down processing is certainly deficient in schizophrenia, future investigations of bottom-up dysfunction will further clarify the underlying causes of cognitive deficits in this disorder.

### 4.4 Implications for future treatment studies

Finally, results reported here also encourage future studies to develop novel intervention strategies, such as complementary sensory-based therapies, which may help to correct early sensory dysfunction and thus further facilitates the remediation of high-order function in schizophrenia. Cognitive training must address limitations in perceptual/pre-attentive processing first (for a review, (Vinogradov et al., 2012)). This hope was preliminarily bolstered by some finding of learning-induced neuroplasticity, in which training in early auditory or visual processes results in substantial gains in verbal or visual cognitive processes through “bottom-up” tuning of the neural systems (Adcock et al., 2009; Biagianti et al., 2016; Dale et al., 2016; Fisher et al., 2009; Hochberger et al., 2019; Surti et al., 2011).

### 4.5 Limitation

Notwithstanding its implications, the main limitations of this study should be acknowledged. A main limitation in the current study, as well as many other clinical imaging studies in the field, is the effect of antipsychotic drugs. While we cannot eliminate completely the potential confounding effects of medication, chlorpromazine equivalents were not associated with the altered gradient or SFC scores. Due to the use of the cross-sectional research design, we did not establish the developmental trajectories of altered cortical hierarchy in schizophrenia. Because altered sensory-motor FC abnormalities have been consistently observed in clinical-high risk, early-stage including drug-naive first-episode, and chronic schizophrenia (Berman et al., 2017; Dong et al., 2018a; Du et al., 2018; Guo et al., 2014; Jiang et al., 2015), it is possible that the prominent compression from the sensorimotor portion of the cortical hierarchy is present at different stages of the illness, possibly ranging from pre-clinical to early and late stages of the disorder. Future longitudinal studies may evaluate the development of cortical hierarchy in schizophrenia across time.

## 4. Conclusions

In conclusion, the present study provided novel system-level substrate underlying the pathological interaction of early sensory and cognitive function in schizophrenia, i.e., the compression of sensorimotor-to-transmodal cortical hierarchy organization. Within the framework of then compressed cortical hierarchy organization, a cascade of impairments stemming from the disrupted somatosensory-motor system and inefficient integration of bottom-up sensory information with attentional demands and executive control processes may partially account for high-level cognitive deficits of schizophrenia. While top-down processing is certainly deficient in schizophrenia, future investigations of bottom-up dysfunction will further clarify the underlying causes of cognitive deficits in this disorder and promote the development of new treatment intervention.

## 5. Acknowledgments

The authors declare no conflicts of interests. This work was supported by the grant from National Key R&D Program of China (2018YFA0701400), The grants from the National Nature Science Foundation of China (grant number: 61933003, 81771822, 81861128001, and 81771925), The Project of Science and Technology Department of Sichuan Province (2019YJ0179), and the CAMS Innovation Fund for Medical Sciences (CIFMS) (No.2019-I2M-5-039).We are grateful to all the participants in this study. Our thanks also go to Dr. Xi Chen (Civil Aviation Flight University of China) and Mr. Xin Chang (University of Electronic Science and Technology of China) for their help to collect the dataset.

