## Supplementary info for "Altered Sensorimotor-to-Transmodal Hierarchical Organization in Schizophrenia": Supplementary Info.docx

**Global signal removal**

Given recent studies found evidence of altered global signal in schizophrenia patients (Hahamy et al., 2014, Yang et al., 2014), supporting the idea that the global signal contains pathophysiologically relevant information, we did not performed global signal regression (GSR) in our main text. However, currently there is no consensus in the neuroimaging field whether to do GSR when computing functional connectivity. To investigate the potential effects of GSR on our findings, we repeated core analyses with GSR, which does not significantly affect trends of overall results (Figure S2-3), although increased SFC degree was found between unimodal seeds and frontoparietal regions, i.e., middle / superior frontal gyrus, inferior parietal lobule, supramarginal gyrus, and dorsal precuneus), and ventral attention regions (dorsal anterior cingulate cortex and bilateral anterior insular cortex / central opercular cortex) at all link-step distances (Step1 to 7). These differences were only found at early and medium link-step distances (Step1 to 4) when without conducting GSR. Overall, the relatively consistent results between without GSR and with GSR indicated the observed main findings reflected the reliable pathophysiologic mechanism of schizophrenia.

**Table S1. Group differences in degree of stepwise functional connectivity**

| **Brain regions** | **T value** | **Voxels (k)** | **MNI coordinates** | | |
| --- | --- | --- | --- | --- | --- |
|  |  |  | **X** | **Y** | **Z** |
| **One step** |  |  |  |  |  |
| ***Patients>Controls*** |  |  |  |  |  |
| L Inferior Temporal Gyrus | 3.90 | 23 | -42 | -54 | -12 |
| L Angular Gyrus / Supramarginal Gyrus / Inferior Parietal Lobule | 4.38 | 49 | -43 | -55 | 55 |
| L Middle Frontal Gyrus | 4.50 | 47 | -36 | 42 | 30 |
| L Superior Frontal Gyrus | 4.92 | 70 | -18 | 12 | 66 |
| Anterior Cingulate Cortex / Supp_Motor_Area | 5.14 | 62 | 0 | 30 | 24 |
| R Middle Frontal Gyrus | 5.17 | 43 | 36 | 36 | 36 |
| L Precuneus Cortex | 5.38 | 50 | -6 | -72 | 54 |
| R Supramarginal Gyrus /Angular Gyrus / Inferior Parietal Lobule | 5.95 | 71 | 54 | -36 | 18 |
| R Insular Cortex / Central Opercular Cortex / Superior Temporal Gyrus | 6.21 | 61 | 48 | 12 | -6 |
| L Temporal Pole / Insular Cortex / Central Opercular Cortex / Superior Temporal Gyrus | 6.35 | 73 | -48 | 6 | -6 |
| ***Patients< Controls*** |  |  |  |  |  |
| R pre/postcentral Gyrus | -7.61 | 200 | 48 | -18 | 42 |
| L pre/postcentral Gyrus | -6.56 | 92 | -54 | -18 | 36 |
| Bilateral Calcarine / Lingual Gyrus / Cuneus | -5.04 | 203 | 6 | -66 | 12 |
| **Two steps** |  |  |  |  |  |
| ***Patients>Controls*** |  |  |  |  |  |
| L Middle Frontal Gyrus | 4.51 | 55 | -36 | 42 | 30 |
| R Middle Frontal Gyrus | 5.18 | 57 | 36 | 36 | 36 |
| L Supramarginal Gyrus / Inferior Parietal Lobule / Angular Gyrus / | 5.48 | 89 | -66 | -48 | 18 |
| L Superior Frontal Gyrus / Anterior Cingulate Cortex / Supp_Motor_Area | 5.60 | 156 | -18 | 12 | 66 |
| R Insular Cortex / Central Opercular Cortex / Superior Temporal Gyrus / Inferior Frontal Gyrus | 5.62 | 68 | 48 | 12 | 0 |
| R Angular Gyrus / Supramarginal Gyrus / Inferior Parietal Lobule | 5.92 | 106 | 54 | -36 | 18 |
| R / L Precuneus Cortex | 6.12 | 148 | 6 | -66 | 66 |
| L Temporal Pole / Insular Cortex / Central Opercular Cortex / Superior Temporal Gyrus / Inferior Frontal Gyrus | 6.63 | 80 | -48 | 6 | -6 |
| ***Patients< Controls*** |  |  |  |  |  |
| R pre/postcentral Gyrus | -7.78 | 260 | 48 | -18 | 42 |
| L pre/postcentral Gyrus | -6.62 | 121 | -54 | -18 | 36 |
| Bilateral Middle Occipital Gyrus / Calcarine / Lingual Gyrus / Cuneus | -5.01 | 424 | 42 | -66 | 0 |
| **Three steps** |  |  |  |  |  |
| ***Patients>Controls*** |  |  |  |  |  |
| L Supramarginal Gyrus / Angular Gyrus / Inferior Parietal Lobule | 4.38 | 80 | -66 | -48 | 18 |
| L Middle Frontal Gyrus | 4.75 | 63 | -36 | 24 | 36 |
| Middle / Inferior Temporal Gyrus / Insular Cortex /Central Opercular Cortex | 5.22 | 84 | 54 | -24 | -18 |
| R Middle Frontal Gyrus | 5.26 | 67 | 36 | 30 | 36 |
| R Angular Gyrus / Supramarginal Gyrus / Inferior Parietal Lobule | 5.72 | 96 | 48 | -48 | 36 |
| L Superior Frontal Gyrus / Anterior Cingulate Cortex / Supp_Motor_Area | 5.74 | 162 | -12 | 18 | 60 |
| R/L Precuneus Cortex / Middle Cingulum Cortex | 6.07 | 124 | 0 | -54 | 72 |
| Insular Cortex / Central Opercular Cortex / Temporal Pole / Middle Temporal Gyrus / Inferior Frontal Gyrus | 6.45 | 90 | -48 | 12 | -6 |
| ***Patients< Controls*** |  |  |  |  |  |
| R pre/postcentral Gyrus | -7.43 | 240 | 48 | -18 | 42 |
| L pre/postcentral Gyrus | -6.46 | 190 | -54 | -18 | 36 |
| Bilateral Middle Occipital Gyrus / Calcarine / Lingual Gyrus / Cuneus | -5.84 | 537 | 42 | -66 | 0 |
| **Four steps** |  |  |  |  |  |
| ***Patients>Controls*** |  |  |  |  |  |
| Middle Cingulum Gyrus | 4.00 | 21 | 0 | -18 | 30 |
| L Middle Frontal Gyrus | 4.27 | 50 | -36 | 12 | 36 |
| R Middle Frontal Gyrus | 4.46 | 53 | 36 | 30 | 36 |
| R Inferior Parietal Lobule / Supramarginal Gyrus /Angular Gyrus | 4.47 | 43 | 42 | -48 | 36 |
| L Supramarginal Gyrus / Inferior Parietal Lobule / Angular Gyrus | 5.13 | 45 | -66 | -48 | 18 |
| Anterior Cingulate Cortex / Superior Frontal Gyrus / Inferior Frontal Gyrus | 5.14 | 113 | -5 | 11 | 58 |
| Inferior / Middle temporal Gyrus / Inferior Frontal Gyrus | 5.42 | 178 | 42 | -6 | -30 |
| Inferior / Middle temporal Gyrus / Inferior Frontal Gyrus | 5.52 | 272 | -46 | -13 | -25 |
| L Insular Cortex/ Central Opercular Cortex | 6.04 | 37 | -48 | 12 | -6 |
| ***Patients< Controls*** |  |  |  |  |  |
| R pre/postcentral Gyrus | -6.84 | 256 | 54 | -12 | 42 |
| L pre/postcentral Gyrus | -6.04 | 170 | -54 | -18 | 36 |
| Bilateral Middle Occipital Gyrus / Calcarine / Lingual Gyrus / Cuneus | -5.83 | 546 | 42 | -66 | 0 |
| **Five steps** |  |  |  |  |  |
| ***Patients>Controls*** |  |  |  |  |  |
| Middle Temporal Gyrus | 4.26 | 20 | 54 | -30 | -12 |
| L Superior Frontal Gyrus / Supp_Motor_Area | 4.93 | 158 | -6 | 12 | 60 |
| L Middle / Inferior Temporal Gyrus / Temporal_Pole / Inferior Frontal Gyrus | 5.19 | 215 | -42 | 6 | -30 |
| Middle / Inferior Temporal Gyrus / Inferior Frontal Gyrus | 5.27 | 101 | 42 | 0 | -36 |
| ***Patients< Controls*** |  |  |  |  |  |
| R pre/postcentral Gyrus | -6.21 | 209 | 54 | -12 | 42 |
| L pre/postcentral Gyrus | -5.95 | 92 | -36 | -30 | 60 |
| Bilateral Middle Occipital Gyrus / Calcarine / Lingual Gyrus / Cuneus | -5.53 | 545 | 42 | -66 | 0 |
| **Six steps** |  |  |  |  |  |
| ***Patients>Controls*** |  |  |  |  |  |
| L Frontal_Sup_Orb | 4.03 | 42 | -18 | 60 | -6 |
| L Superior Frontal Gyrus / Supp_Motor_Area | 4.51 | 84 | -6 | 12 | 60 |
| R Middle Temporal Gyrus | 4.84 | 29 | 60 | -36 | -6 |
| L Middle / Inferior Temporal Gyrus /Temporal_Pole / Inferior Frontal Gyrus | 5.60 | 212 | -42 | 6 | -30 |
| R Middle / Inferior Temporal Gyrus / Temporal_Pole | 5.65 | 109 | 42 | 0 | -36 |
| ***Patients< Controls*** |  |  |  |  |  |
| R pre/postcentral Gyrus | -5.92 | 227 | 48 | -18 | 42 |
| L pre/postcentral Gyrus | -5.50 | 180 | -36 | -30 | 60 |
| Bilateral Middle Occipital Gyrus / Calcarine / Lingual Gyrus / Cuneus | -5.26 | 512 | 42 | -66 | 0 |
| **Seven steps** |  |  |  |  |  |
| ***Patients>Controls*** |  |  |  |  |  |
| L Middle Frontal Gyrus | 3.44 | 22 | -24 | 54 | 18 |
| Superior-Medial Frontal Gyrus | 4.25 | 21 | 0 | 42 | 42 |
| L Superior Frontal Gyrus / Supp_Motor_Area | 4.27 | 42 | -6 | 12 | 60 |
| R Middle Temporal Gyrus | 4.84 | 31 | 60 | -36 | -6 |
| R Middle / Inferior Temporal Gyrus /Temporal_Pole | 5.78 | 119 | 42 | 0 | -36 |
| L Middle / Inferior Temporal Gyrus / Temporal_Pole / Inferior Frontal Gyrus | 5.92 | 220 | -42 | 6 | -30 |
| ***Patients< Controls*** |  |  |  |  |  |
| R pre/postcentral Gyrus | -5.77 | 200 | 48 | -18 | 42 |
| L pre/postcentral Gyrus | -5.27 | 196 | -36 | -30 | 60 |
| Bilateral Middle Occipital Gyrus / Calcarine / Lingual Gyrus / Cuneus | -5.26 | 486 | 42 | -66 | 0 |

**Notes:** L, left side of brain; R, right side of brain. Results are reported using a voxel-wise FDR threshold of *P* < .05 and an additional cluster-size threshold of k=20.

Table S2. Association Between Atypical Gradient, SFC Degree and Clinical Severity in Schizophrenia

| Index | Region | | PANSS-P | | PANSS-N | | PANSS-G | | PANSS-T | |
| --- | --- | --- | --- | --- | --- | --- | --- | --- | --- | --- |
|  |  |  | r | p | r | p | r | p | r | p |
| Gradient | Ventral Medial Frontal Gyrus | | -0.032 | 0.799 | -0.318 | 0.010 | -0.031 | 0.810 | -0.173 | 0.172 |
| Gradient | L Anterior Insula | | -0.007 | 0.953 | -0.258 | 0.039 | 0.092 | 0.470 | -0.155 | 0.222 |
| Gradient | L Precuneus | | -0.366 | 0.002 | -0.102 | 0.423 | -0.183 | 0.147 | -0.292 | 0.019 |
| SFC-step1 | L Superior Frontal Gyrus | | -0.085 | 0.499 | -0.051 | 0.688 | -0.278 | 0.026 | -0.184 | 0.144 |
|  | R Anterior Insular Cortex / Central Opercular Cortex | | -0.290 | 0.020 | -0.178 | 0.157 | -0.303 | 0.014 | -0.345 | 0.005 |
|  | L Anterior Insular Cortex / Central Opercular Cortex | | -0.354 | 0.004 | -0.221 | 0.078 | -0.259 | 0.038 | -0.374 | 0.002 |
|  | R pre/postcentral Gyrus | | 0.250 | 0.046 | 0.090 | 0.476 | 0.353 | 0.004 | 0.309 | 0.012 |
|  | L pre/postcentral Gyrus | | 0.186 | 0.139 | 0.152 | 0.227 | 0.310 | 0.012 | 0.290 | 0.020 |
|  | R Lingual Gyrus / Cuneus | | 0.324 | 0.008 | -0.104 | 0.389 | -0.036 | 0.776 | 0.079 | 0.531 |
| SFC-step2 | L Anterior Insular Cortex / Central Opercular Cortex | | -0.238 | 0.057 | -0.343 | 0.005 | -0.276 | 0.027 | -0.386 | 0.001 |
|  | R pre/postcentral Gyrus | | 0.243 | 0.052 | 0.163 | 0.197 | 0.379 | 0.002 | 0.351 | 0.004 |
| SFC-step3 | L Anterior Insular Cortex / Central Opercular Cortex | | -0.159 | 0.208 | -0.383 | 0.001 | -0.252 | 0.044 | -0.358 | 0.003 |
|  | R pre/postcentral Gyrus | | 0.164 | 0.194 | 0.166 | 0.190 | 0.326 | 0.008 | 0.293 | 0.018 |
|  | R Middle Occipital Gyrus / Lingual Gyrus / Cuneus | | 0.166 | 0.190 | 0.299 | 0.016 | 0.026 | 0.833 | -0.050 | 0.693 |
| SFC-step4 | L Middle Frontal Gyrus | | 0.050 | 0.689 | 0.334 | 0.006 | 0.102 | 0.418 | 0.220 | 0.079 |
|  | L Anterior Insular Cortex / Central Opercular Cortex | | -0.025 | 0.839 | -0.369 | 0.002 | -0.226 | 0.072 | -0.280 | 0.024 |
|  | R Middle Occipital Gyrus / Lingual Gyrus / Cuneus | | 0.163 | 0.198 | 0.293 | 0.018 | 0.015 | 0.904 | -0.054 | 0.671 |
| SFC-step5 | L pre/postcentral Gyrus | | 0.029 | 0.816 | 0.324 | 0.008 | 0.247 | 0.048 | 0.270 | 0.030 |
|  | R Middle Occipital Gyrus / Lingual Gyrus / Cuneus | | 0.109 | 0.389 | 0.247 | 0.049 | -0.024 | 0.850 | -0.074 | 0.559 |
| SFC-step6 | R pre/postcentral Gyrus | | 0.049 | 0.696 | 0.170 | 0.177 | 0.262 | 0.036 | 0.215 | 0.086 |
|  | L pre/postcentral Gyrus | | 0.010 | 0.933 | 0.313 | 0.011 | 0.255 | 0.041 | 0.260 | 0.037 |
|  | R Middle Occipital Gyrus / Lingual Gyrus / Cuneus | | 0.066 | 0.601 | 0.259 | 0.038 | -0.069 | 0.587 | -0.118 | 0.349 |
| SFC-step7 | R pre/postcentral Gyrus | | 0.039 | 0.753 | 0.169 | 0.181 | 0.255 | 0.041 | 0.207 | 0.09 |
|  | L pre/postcentral Gyrus | | 0.006 | 0.961 | 0.307 | 0.013 | 0.256 | 0.041 | 0.256 | 0.041 |
|  | R Middle Occipital Gyrus / Lingual Gyrus / Cuneus | | 0.057 | 0.648 | 0.264 | 0.035 | -0.073 | 0.538 | -0.128 | 0.309 |
|  |  | SZ>HC | Higher value is associated to worse clinical symptoms | | | | | | | |
|  |  | SZ<HC | Higher value is associated to better clinical symptoms | | | | | | | |

Notes: L, left side of brain; R, right side of brain. PANSS-P, PANSS-Positive Symptoms; PANSS-N, PANSS-Negative Symptoms; PANSS-G, PANSS-General Symptoms; PANSS-T, PANSS-Total Symptoms. Note that higher scores in PANSS indicate increased severity of symptoms.

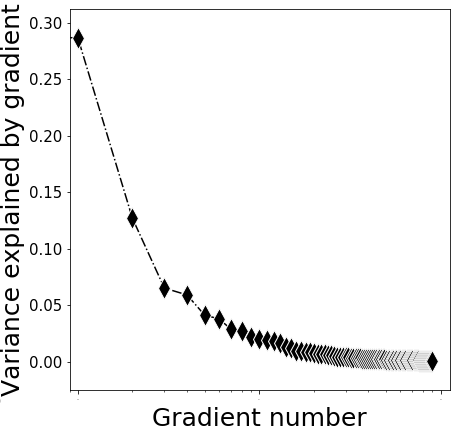

Figure S1. Variance explained by gradient.

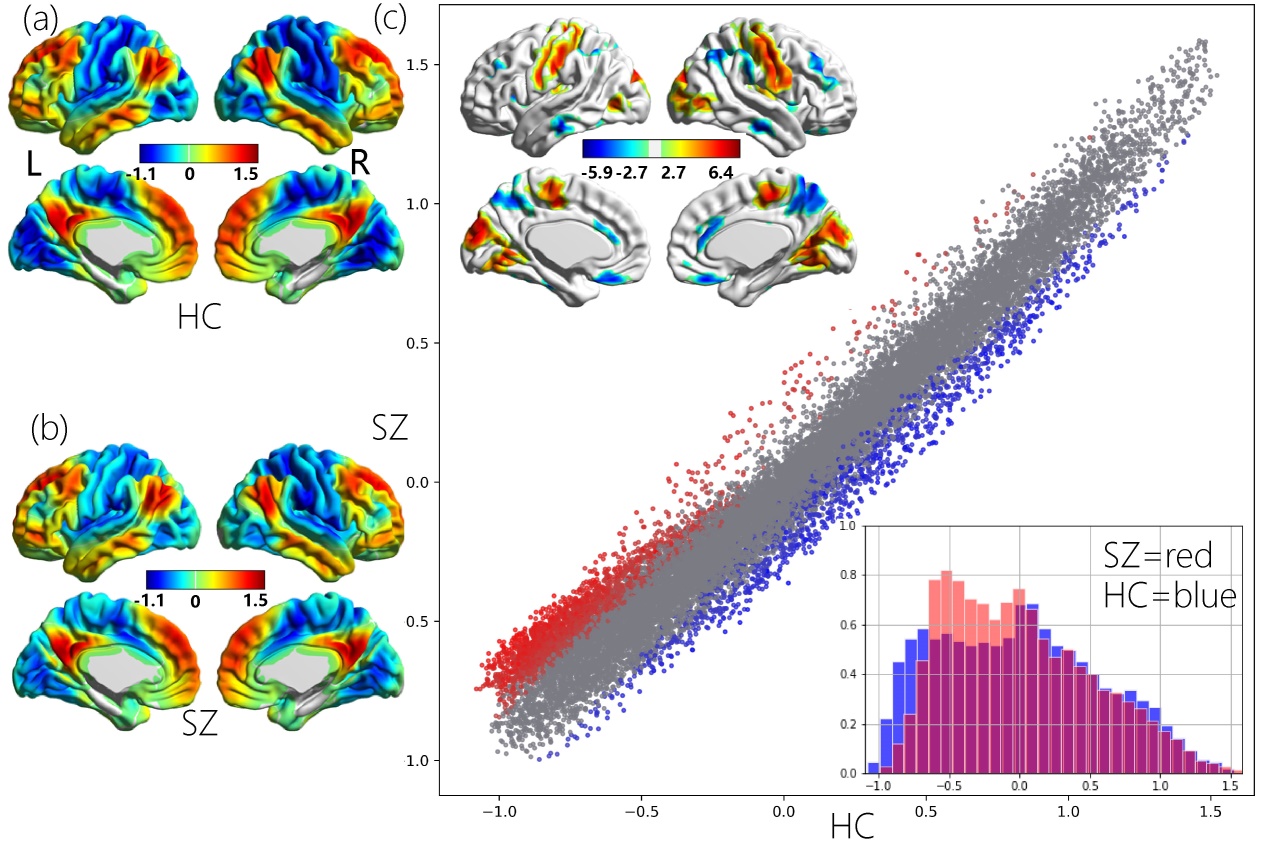

Figure S2. Group patterns and differences in the cerebral principal functional gradient with GSR. (a) Gradient pattern in HC. (b) Gradient pattern in SZ. (c) Group differences between SZ and HC. Scatterplot represents cerebral gradient of SZ (y axis) vs. cerebral gradient of HC (x axis). Scatterplot colors correspond to group differences map as shown in top left corner of Figure S2(c): higher gradient value in SZ (red), and lower gradient value in SZ (blue) compared to HC. Compressed gradient pattern in SZ is shown in density histograms in bottom right corner of Figure S2(c) All results are shown after FDR correction (P < 0.05).

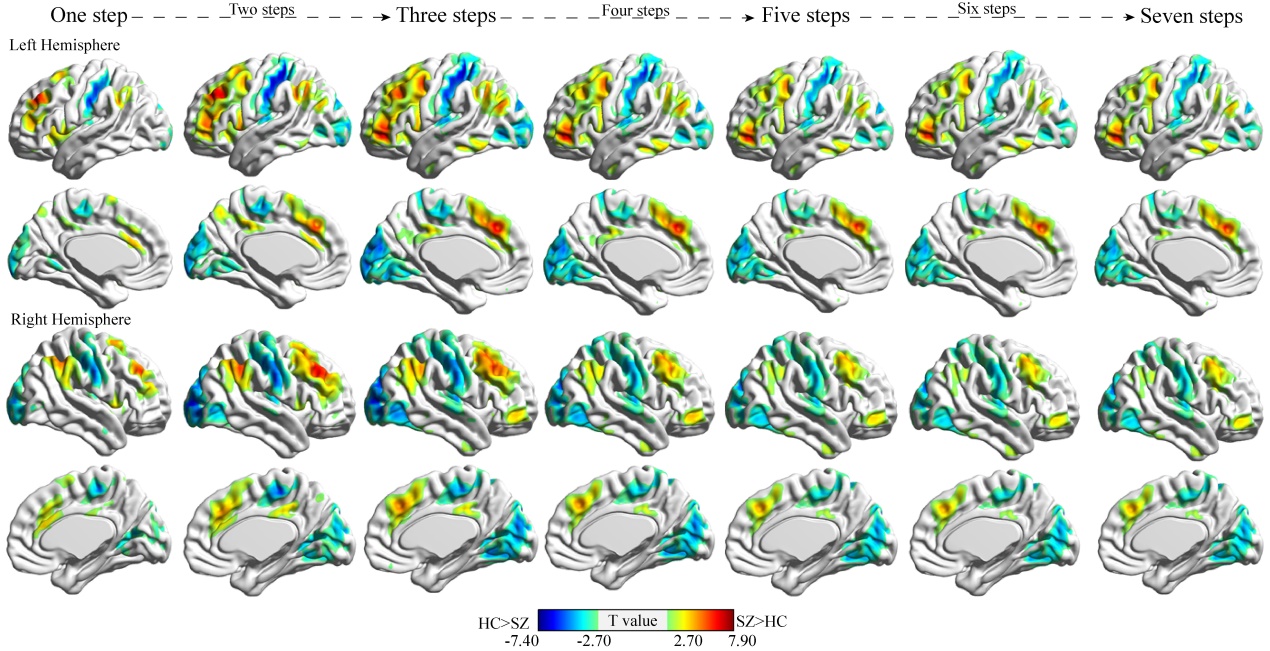

Figure S3. Group differences between schizophrenia and healthy control (HC) in stepwise functional connectivity degree with GSR. All results are shown after FDR correction (P < 0.05).
